# Studying the precuneus reveals structure-function-affect correlation in long-term meditators

**DOI:** 10.1101/822056

**Authors:** Aviva Berkovich-Ohana, Edna Furman-Haran, Rafael Malach, Amos Arieli, Michal Harel, Sharon Gilaie-Dotan

## Abstract

Understanding the relationship between brain structure, function and self-reports has hardly been addressed until now in meditation research. Here we demonstrate such relationship, using Mindfulness meditation (MM). MM aims to reduce thought-related processes and enhance bodily awareness, thereby reducing identification with thought content and deconstructing maladaptive self-schema. We thus hypothesized that structure of the default mode network (DMN) regions, associated with spontaneous thoughts and self-representation, would negatively correlate with MM experience and self-reported positive affect, while positively correlating with DMN resting-state function.

Cross-sectionally comparing a unique group of adept MM practitioners and meditation-naïve matched controls using voxel based morphometry revealed that gray matter (GM) density of the left precuneus (L-Prc) was negatively correlated with MM expertise. Furthermore, GM density of the L-Prc was positively correlated with resting state and task related functional (fMRI) measures within the L-Prc in the MM practitioners, but not in the controls. Finally, the L-Prc’s GM density negatively correlated with positive affect across all participants.

Our findings may shed light on understanding structure-function-self reports relationship. While our approach enables studying suggestive correlations in expert MM practitioners, longitudinal studies are required for direct insights concerning the question of causality.

**Significance statement:** Understanding the relationship between brain structure and function, individual differences and self-reports, is an important goal of neuroscience. Yet, these factors were hardly investigated together in meditation research. The precuneus is part of the default mode network (DMN) involved in thought related processes. Mindfulness meditation (MM) is a mental practice aiming to reduce thought related processes. Here, by cross-sectionally comparing adept meditators and controls, we found that in the precuneus of MM practitioners, structure associated with function. Overall, structure negatively correlated with practice length, as well as with positive affect scores. The structure-function correlation was only significant in the meditators group, possibly implying that prolonged meditation improved structure and function attunement in the DMN. More generally, this study demonstrates that mental practice can be related to conjoint structural and functional effects, as well as to affective self-reports.

## Introduction

Understanding the relationship between the structure and function in the human brain, and how they are associated with individual differences, is one of the most important goals of neuroscience. Inter-individual differences are evident in behavior (e.g. Greenwald, McGhee, & Schwartz, 1998; Raz et al., 2005), but whether these behavioral differences are manifested in both structure and function remains unclear. Up until now, inter-individual differences studies have addressed links between behavior and function (e.g. Gilaie-Dotan et al., 2013; Hubbard, Arman, Ramachandran, & Boynton, 2005) or behavior and structure (e.g. Fleming, Weil, Nagy, Dolan, & Rees, 2010; Schwarzkopf, Song, & Rees, 2011), but hardly addressed the link between all of these three factors together in general, and specifically in meditation research (but see recent paper by Yang, Barrós-Loscertales & Li et al., 2019). This study aims at filling this gap.

Meditation is rapidly spreading as a secular practice worldwide, with tens of millions performing daily practice in the USA alone, and with accelerating increase in scientific publications (Davidson & Kaszniak, 2015; Tang, Hölzel, & Posner, 2015). Meditation training-induced individual differences are well acknowledged in psychological studies of behavior (e.g. Bornemann, Herbert, Mehling, & Singer, 2015; Brown, Forte, & Dysart, 1984; Jha, Krompinger, & Baime, 2007), self-reports (e.g. Baer, Smith, Hopkins, Krietemeyer, & Toney, 2006; Barbosa et al., 2013; Berkovich-Ohana & Glicksohn, 2015) including phenomenology (Abdoun, Zorn, Poletti, Fucci, & Lutz, 2019; Petitmengin, van Beek, Bitbol, Nissou, & Roepstorff, 2018; Przyrembel & Singer, 2018), or meditation expertise (Cahn & Polich, 2009; Chan & Woollacott, 2007; recently reviewed by Falcone & Jerram, 2018; Fox et al., 2014). Meditation-related effects have also raised much interest in the last decade in functional neuroimaging studies (e.g. Berkovich-Ohana, Harel, Hahamy, Arieli, & Malach, 2016; Brewer et al., 2011; Pagnoni, 2012; Tang et al., 2007) as well as structural studies (e.g. Fox et al., 2014; Hölzel, Carmody, et al., 2011; Lazar et al., 2005). However, linking meditation-related individual differences in structure, function, training expertise and self-reported experience was not previously reported, to the best of our knowledge. Here, we report such an intricate structure – function – self-reports – training expertise relationship, within the context of mindfulness meditation.

Mindfulness meditation (MM) is a non-sectarian Western development of the Buddhist Theravada Vippassana meditation, aiming at focusing non-judgmental awareness to momentary experience (Baer, 2003; Kabat-Zinn, 2011). MM is often referred to as “Insight Meditation”, as the purpose of this practice is to advance insight concerning the nature of reality, including especially the lack of any persistent personal self (Dreyfus & Thompson, 2007; Gunaratana & Gunaratana, 2011). Surging research shows that MM generally exerts beneficial effects on physical and mental health, including heightened attention and emotion regulation (Tang et al., 2015), increased immune function (Black & Slavich, 2016), and possibly offsetting age-related cognitive decline (Gard, Hölzel, & Lazar, 2014).

Notably, one of the proposed key mechanisms for the effects of mindfulness is reducing identification with a rigid self-concept through enhanced meta-awareness (Dambrun & Ricard, 2011; Hart, 1987; Olendzki, 2010), which creates a shift in self-awareness and its proposed underling neural activity (Dahl, Lutz, & Davidson, 2015; Dor-Ziderman, Berkovich-Ohana, Glicksohn, & Goldstein, 2013; Hölzel, Lazar, et al., 2011; Tang et al., 2015; Vago & Silbersweig, 2012), or as phrased by Tang et al. (2015, 2019): “According to Buddhist philosophy, the identification with a static concept of ‘self’ causes psychological distress. Dis-identification from such a static self-concept results in the freedom to experience a more genuine way of being. Through enhanced meta-awareness (making awareness itself an object of attention), mindfulness meditation is thought to facilitate a detachment from identification with the self as a static entity and a tendency to identify with the phenomenon of ‘experiencing’ itself is said to emerge.” Specifically, it is conceptualized as enhancing regulation of attention and meta-awareness, which in turn down-regulates automatic process of absorption in the contents of consciousness (experiential-fusion), as well as deconstructing mal-adaptive self-schema by employing self-inquiry to foster insight into self-related psychological processes (Dahl, Lutz, & Davidson, 2015).

The cortical network largely accepted to be involved in self-referential processing is the default mode network (DMN, Buckner, Andrews-Hanna, & Schacter, 2008; Raichle et al., 2001), which was shown to support autobiographic memory and mental time traveling to past – i.e. episodic memories - or future, i.e. planning (Addis, Wong, & Schacter, 2007; Bar, 2007; Christoff, Cosmelli, Legrand, & Thompson, 2011; Gusnard, Akbudak, Shulman, & Raichle, 2001; Kim, 2012; Northoff et al., 2006; Schacter, Addis, & Buckner, 2007). The DMN habitually activates during rest and mind-wandering and deactivates during an external task (Mason et al., 2007; Smallwood & Schooler, 2006). It classically includes the medial prefrontal cortex, inferior parietal lobule (IPL), posterior cingulate cortex / precuneus (PCC/Prc), and the medial temporal lobe (Buckner et al., 2008; Raichle et al., 2001).

Accumulating findings using blood oxygenated level dependent (BOLD)-fMRI demonstrate that one of the major effects of MM practice are connectivity and functional alterations associated with the DMN. Reports of such resting-state alterations in functional connectivity in meditators include both reduced (Berkovich-Ohana et al., 2016; Garrison et al., 2013; Taylor et al., 2013) and increased (Jang et al., 2011; Taylor et al., 2013; Yang et al., 2016) functional connectivity among various DMN nodes, as well as altered functional connectivity between DMN nodes and regions outside of the DMN including the cognitive control network (Brewer et al., 2011; Creswell et al., 2016), sensory regions (Farb et al., 2007; Froelinger, 2012; Josipovic, 2012; Kilpatrick et al., 2011), subcortical regions (Shao et al., 2016), and the orbitofrontal cortex (Hasenkamp & Barsalou, 2012; Jang et al., 2011). More relevant to our study, direct measurement of BOLD fMRI activity during mindfulness-related practices revealed that regions of the DMN show relatively little activity in meditators compared to novice control participants (Brewer et al., 2011; Farb et al., 2007; meta-analysis by Fox et al., 2016; Ives-Deliperi, Solms, & Meintjes, 2011; Pagnoni, Cekic, & Guo, 2008; Pagnoni, 2012), but see also opposite effects have been reported (Hölzel et al., 2007; Xu et al., 2014). This was largely interpreted as indicating diminished self-referential processing during MM. Recently, these MM state findings of reduced DMN activity during meditation were extended beyond the meditative state *per se* (Berkovich-Ohana, Harel, Hahamy, et al., 2016; Garrison, Zeffiro, Scheinost, Constable, & Brewer, 2015). Specifically, we showed that relatively lower DMN activity – specifically Prc/PCC in adept meditators compared to controls is also found during a visuo-attentional task and during resting-state, rendering this effect a trait (long-term) condition (Berkovich-Ohana, Harel, Hahamy-Dubossarsky, Arieli, & Malach, 2016). As the posterior node of the DMN, the PCC, was related using real-time fMRI neurofeedback to ‘identifying with attributes of ourselves’, or ‘being caught up in experience’ (Brewer & Garrison, 2014; Brewer, Garrison, & Whitfield-Gabrieli, 2013; Garrison et al., 2013), the findings of reduced DMN activity in MM practitioners as a state and trait effect lend support to the theory that mindfulness achieves its positive outcomes through a process of dis-identification from the content of ones’ thoughts (Dahl et al., 2015; Dambrun & Ricard, 2011; Hadash, Plonsker, Vago, & Bernstein, 2016; Hadash, Segev, Tanay, Goldstein, & Bernstein, 2016; Vago & Silbersweig, 2012).

While fMRI studies largely show that MM practice is associated with reduced DMN activity, the related structural effects in the DMN are less clear (recently reviewed by Fox et al., 2014). Specifically, several neuroanatomical studies investigated PCC gray matter (GM) thickness: while one study indicated reduction in meditators relative to controls (Kang et al., 2013), two other studies reported GM increases following a short 8 weeks MM intervention (Hölzel, Carmody, et al., 2011; Yang et al., 2019), and another study failed to find group differences (Grant et al., 2013).

In this study we investigated a unique group of adept MM practitioners and a group of matched controls, for which we recently reported reduced DMN BOLD-fMRI activity during resting-state and task for the MM group compared to the control group (Berkovich-Ohana, Harel, Hahamy, et al., 2016; Berkovich-Ohana, Harel, Hahamy-Dubossarsky, et al., 2016). Here, we examined GM density individual differences, and their possible modulation by MM training expertise. Additionally, we explored whether these neuroanatomical differences are associated with function (fMRI activity) or self-reported measures of undistracted awareness and contentment.

## Methods

### Participants

The study design was cross-sectional: Eighteen healthy mindfulness meditators (average age ± standard deviation (SD): 43.7 ± 10.6 years, 6 females) and eighteen meditation-naïve participants that served as controls (average age ± SD 42.4 ± 9.7 years, 4 females) participated in the study. This sample size (*n* = 38 total number) marginally enables within-subject correlations between the variables, based on a power analysis which yielded a sample size of *n* = 37 to achieve a power = .80 for one tailed test at alpha = .05. The meditators were long-term practitioners (average 7,560 hours; range 940 – 29,300 hours of formal practice, 15.5 ± 6.7 years), all practicing meditation according to the Satipathana and Theravada Vipassana traditions, and were recruited via the Israeli Insight Society TOVANA. The controls were matched for age, race (all Caucasian), and education (all having university level education). All participants were right handed by self-report, had no history of neurological disorders, and provided written informed consent to participate in the study. High-resolution anatomical MRI scans were obtained for all participants. Out of these 36 participants, only 33 participants (16 MM and 17 controls) underwent fMRI scanning (participated in the structure-function correlation analyses) and thus only those are reported here for structure-function correlations. One MM participant did not fill out the personal report measures, and was thus excluded from structure-self report correlation analyses. The study and experimental procedures were approved by the Tel Aviv Sourasky Medical Center Helsinki committee.

### Self-reported measures

To estimate the participant’s experiences of undistracted awareness we used the Tellegen Absorption Scale (TAS, well-accepted in the field of personality studies) taken from the Multidimensional Personality Questionnaire (Hebrew version from Glicksohn, 1991; Tellegen & Atkinson, 1974). Absorption is a personality trait related to hypnotic ability, which involves the ability to highly focus attention, as well as “openness to absorption and self-altering experiences” (Tellegen & Atkinson, 1974, p. 274). A core feature of absorption is an experience of focused attention wherein: “*absorbed attention amplifies greatly the experience of one part of reality, while other aspects recede from awareness. Consequently, the vivid subjective reality experienced during episodes of absorbed attention may well, in retrospect, during more…normal states of wakefulness, impress one as ‘altered’*” (Tellegen & Atkinson, 1974, p. 274). What becomes altered is the regular sense of self “*objects of absorbed attention acquire an importance and intimacy that are normally reserved for the self and may, therefore, acquire a temporary self-like quality*” (Tellegen & Atkinson, 1974, p. 275). The TAS consists of 34 items, which participants score as true or false (highest score is 34, signifying a high trait ability to experience an altered sense of self), including questions like: “While watching a show, or a play, I may become so involved that I forget about myself and my surrounding and experience the story as if it were real and as if I were taking part in it”, and “Sometimes I ‘step outside’ my usual self and experience an entirely different state of mind.”

To assess contentment, we used the positive affect subscale from the Positive Affect and Negative Affect Schedule (PANAS) (Hebrew version taken from Binenboim, 2003; Watson, Clark, & Tellegen, 1988). The PANAS includes two 10-item mood sub-scales, positive and negative, designed to be taken both in state and trait forms. The 10 descriptors for the positive-affect (PA) scale are: attentive, interested, alert, excited, enthusiastic, inspired, proud, determined, strong and active; and the 10 descriptors for the negative-affect (NA) scale include: distressed, upset, hostile, irritable-angry, scared, afraid-fearful, ashamed, guilty, nervous, and jittery. We used the trait form, asking participants to report to what extent did they generally (in the last year) feel each descriptor. Participants answered on a five-point Likert scale, including: 1 (very slightly or not at all), 2 (a little), 3 (moderately), 4 (quite a bit), and 5 (extremely), highest scores on each subscale being 50.

### Imaging setup

Images were acquired on a 3 Tesla Trio Magnetom Siemens scanner, equipped with a 12-channels head matrix coil (Siemens, Erlangen Germany), located at the Weizmann Institute of Science, Rehovot, Israel. Foam cushions were used for head stabilization, and MR compatible earphones (MR Confon, Magdeburg, Germany) for reducing external noise. High-resolution T1-weighted anatomical images were acquired (1×1×1 mm^3^, 3D MPRAGE, TR=2300 ms, TE=2.98 ms, TI=900ms, flip angle=9°) and were used in the structural analysis and to facilitate the incorporation of the functional data into 3D space. Functional T2*-weighted images were obtained with gradient-echo echo planar imaging sequence (46 axial slices, 3mm thickness without gaps, TR=3000 ms, TE=30 ms, flip angle=90°, FOV=240 mm, matrix size=80×80, resulting in 3×3×3 mm^3^ voxels covering the whole brain).

### Voxel-based morphometric analysis

#### Preprocessing

Voxel-based morphometric analysis was performed with SPM8 (http://www.fil.ion.ucl.ac.uk/spm). For each participant the T1-weighted images were first converted to nifti format with MRIconvert version 2.0 (http://lcni.uoregon.edu). Then, the images were segmented to gray matter, white matter (WM) and cerebrospinal fluid, using the segmentation tools in SPM8. This was followed by diffeomorphic anatomical registration through exponentiated Lie Algebra (DARTEL) for inter-participant registration of the GM images (Ashburner, 2007). The registered images were smoothed with a Gaussian kernel (FWHM = 8 mm) and were then transformed to MNI stereotactic space using affine and non-linear spatial normalization implemented in SPM8 for multiple regression analysis while preserving the amount of signal (i.e. without modulation).

#### Voxel-based morphometry (VBM) second-level modeling and analyses

The following typical VBM analyses were performed in SPM8 (see Results and Figure 1). In line with our hypothesis, we started by examining anatomical correlates of meditation experience in the DMN network, and for that purpose we ran standard VBM analysis in predefined ROIs of the DMN (see ‘Regions of Interest (ROI) definitions’ section below). These hypothesis driven analyses included a group comparison to identify any group differences. We adopted a multiple regression model with binary predictor (meditation yes/no). We also ran an additional multiple regression analysis to identify any correlation of GM density with meditation experience (years of meditation). Age, gender and the total GM density were included in each of these VBM analyses (i.e. in their design matrices) as covariates of no interest to regress out any effects attributable to them. To complement the hypothesis driven analyses, we also performed whole-brain analyses with these same design matrices, but this time not confining the search space to our hypothesis: *F* contrast were computed with *p* = 0.001 (uncorrected) and minimum cluster size=40 voxels were chosen as the criterion to detect whole‐brain voxels with significant effects.

**Figure 1.**
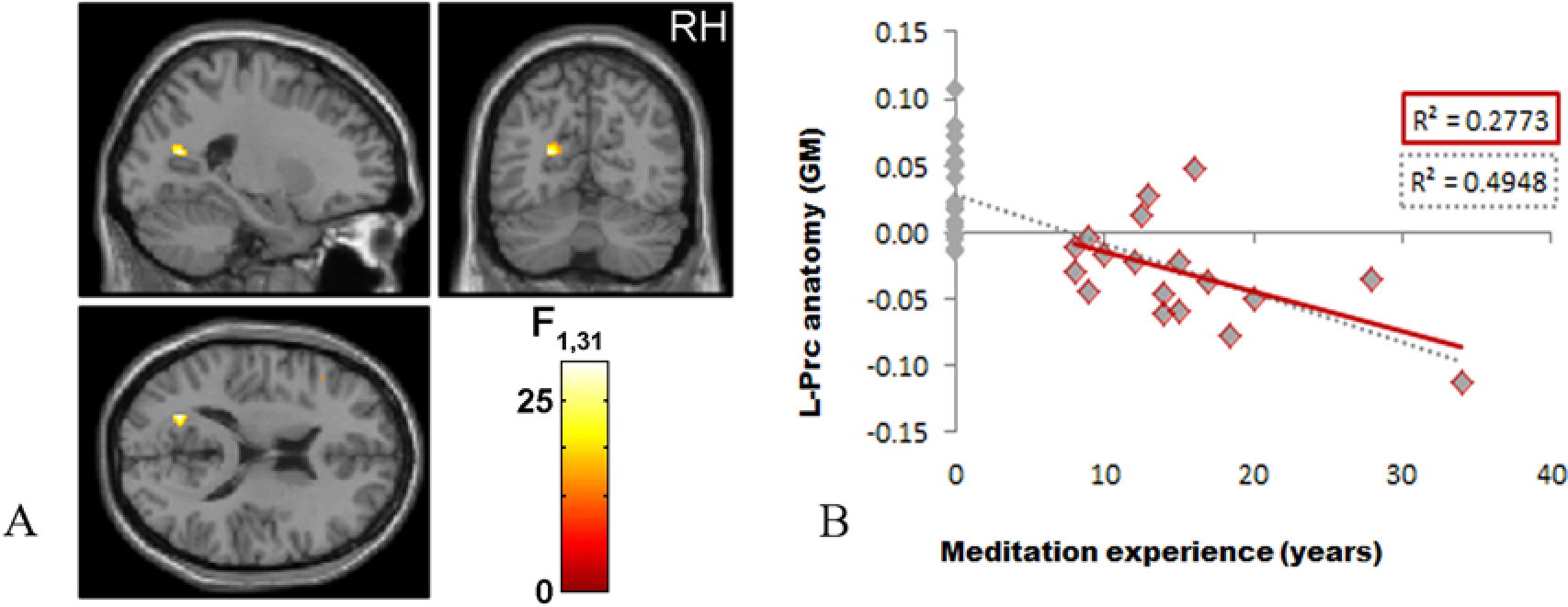
Neuroanatomy of L-Prc is associated with meditation experience. **A.** Second-level VBM analyses found a significant between-group GM thickness difference in the L-Prc (at MNI coordinates −21, −64, 16, *p* =.013, FWE corrected, small volume correction) but not in the other DMN ROIs. The scale bar denotes the *F* statistics (*n* = 36). **B.** To demonstrate that the observed correlations were not driven by outliers, for each participant (*n* = 36) L-Prc peak GM density (in normalized arbitrary units, y axis) is plotted in gray against meditation expertise (in years), the regression line in gray (please note that the y-axis shows normalized values – i.e. in arbitrary units, as extracted from the peak SPM result presented in A). Note that this should not be used for inference as it is not independent of the VBM analysis in (A) and is presented for visualization purposes only. The correlation based on the MM group alone (n=18) that is indicated by the dots with red border and by the red regression line is also significant (R = −0.52661948, t(16)=−2.478, p=0.025). As can be seen the slope is similar in both analyses. RH, Right hemisphere.

#### Regions of interest (ROI) definitions

We focused our second level VBM standard analyses (see above) on four DMN regions - the bilateral precuneus (Prc) and inferior parietal lobule (IPL). These regions of interest (ROIs: L-Prc, R-Prc, L-IPL, R-IPL) were selected based on the group functional fMRI results in the same cohort (Berkovich-Ohana, Harel, Hahamy, et al., 2016). Based on functional criteria, we used the negative group-level activation peaks (based on *n* = 33: 17 MM and 16 control participants) in the visual localizer as the centers of the DMN ROIs. The coordinates of these four negative peaks were converted from Talairach space (L-Prc: −8 −53 26; R-Prc: 3 −54 23; L-IPL: −47 −68 24; R-IPL: 44 −60 25) to MNI space (Laird et al., 2010) using Tal2MNI MATLAB function (MNI coordinates: L-Prc: −8 −56 25; R-Prc: 3 −56 22; L-IPL: −47 −71 22; R-IPL: 44 −63 24). The ROIs were defined as spheres of 10 mm radius around each of these four negative peaks, and were used for small volume correction (SMC) for the ROI-based second level VBM analysis (see above and Figure 1), and for GM density sampling to be compared with functional or affect scores and statistical analyses (see the following section and Figures 2 and 3).

**Figure 2.**
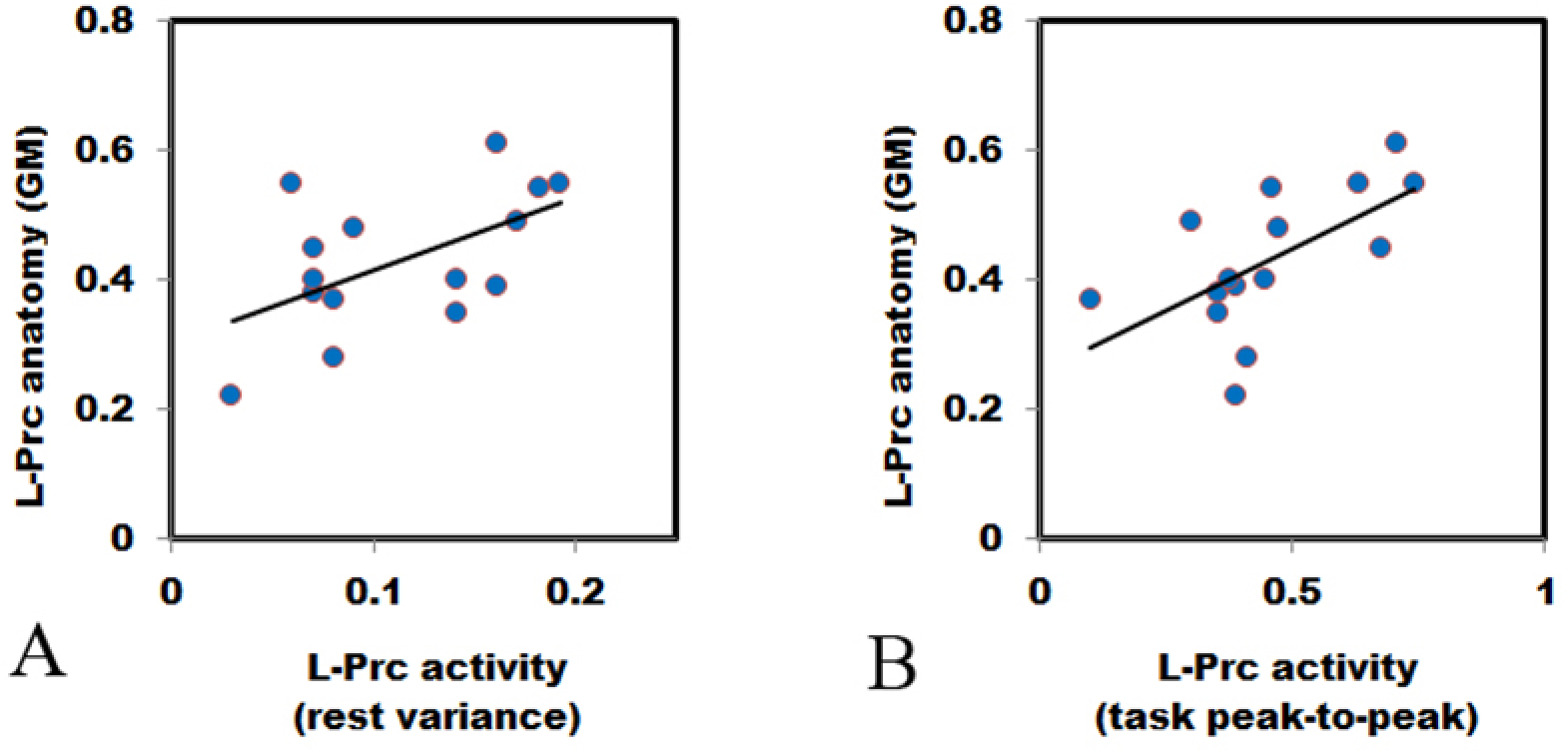
L-Prc structure-function correlation in the MM group. L-Prc GM (y-axis) of the MM group (*n* = 15) is significantly correlated with (A) resting-state variance (x-axis, r = .552, t (13) = 2.39, p = .033) and with (B) task peak-to-peak amplitudes (x-axis, r = .605, t (13) = 2.74, p = .017). The extracted L-Prc ROI GM density (see Methods) is independent of the acquired functional measures. Each point represents an individual MM participant. No significant correlations were found in the control group (Table 1).

**Table 1.** Pearson correlations between L-Prc ROI neuroanatomical gray matter (GM) density (structure) and fMRI BOLD functional measures (*Z*-transformed variance during resting-state, and peak-to-peak activation values during a visual task) or self-reported scores on questionnaires (PANAS and TAS – see Methods). MM – Mindfulness meditators (*n* = 16); Controls (*n* = 17); * *p* < .05; ** *p* < .005 (exact p-values are shown to enable comparisons with Holm-Bonferroni correction).

| Correlated variable |  | Correlation coefficients (*p*-value) |  |  |
| --- | --- | --- | --- | --- |
|  |  | MM | Control | All |
| **fMRI**<br>**measure** | Rest variance | .552* (0.033) | -.102 (ns) | .270 (ns) |
|  | Task peak-to-peak | .605* (0.017) | -.025 (ns) | .294 (ns) |
| **Self-**<br>**reported**<br>**measures** | Positive Affect (PA) | -.555* (0.033) | -.371(ns) | -.522**<br>(0.002) |
|  | Negative Affect (NA) | .028 (ns) | .425 (ns) | .158 (ns) |
|  | TAS | -.426 (ns) | .169 (ns) | -.305 (ns) |

**Figure 3.**
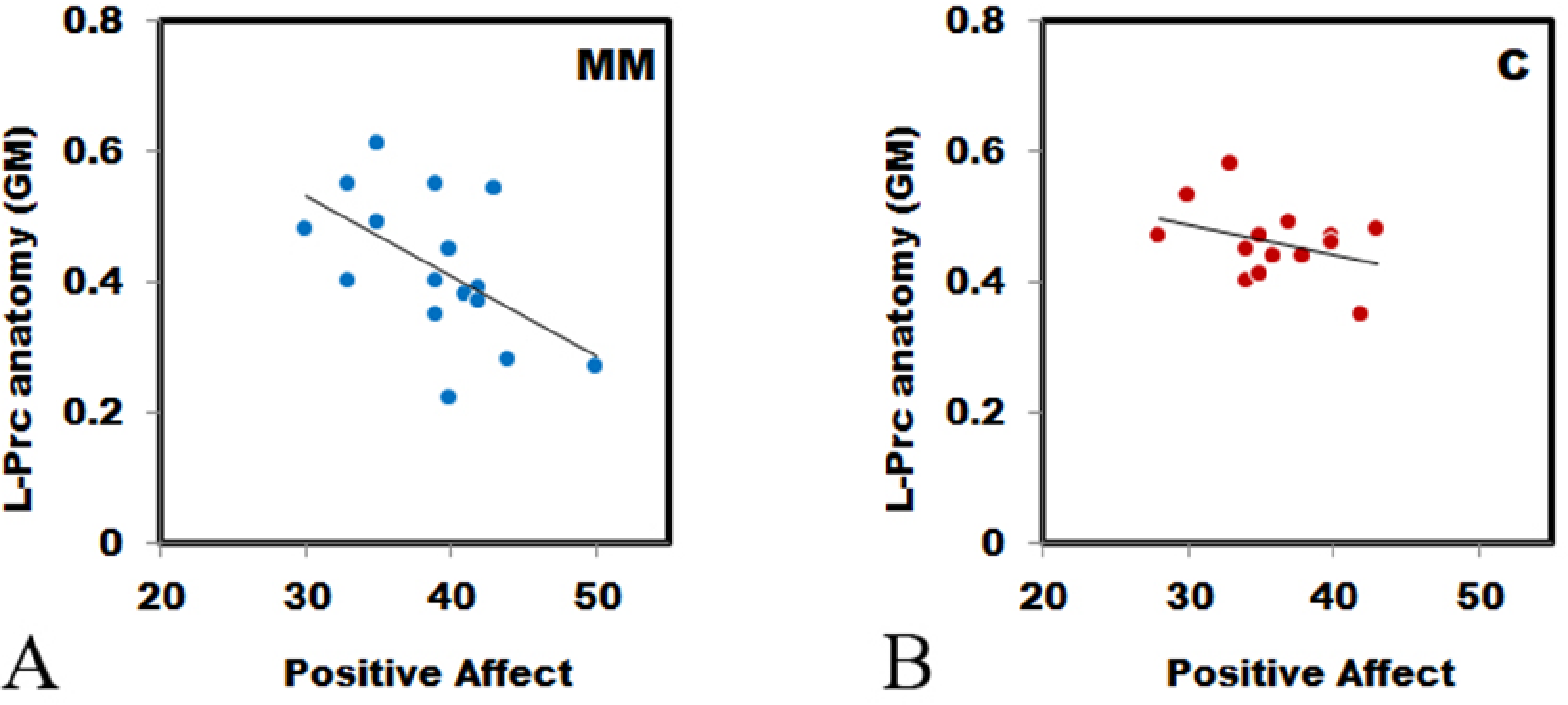
L-Prc neuroanatomy correlates with positive affect. Correlation between L-Prc ROI extracted gray matter (GM) density (y-axis) and positive affect (as measured by the PANAS, x-axis) is significant for the MM group ((A), *n* = 16, *r* = −.555, *p* = .026), but not for the control group ((B), *n* =17, *r* = −.371, *p* = .143), see Methods for more details.

#### GM density sampling

In addition to the VBM second-level analyses, we also performed individual based analysis. To that end we extracted individual GM density from each of the ROIs to be used for between group analyses and to compare with the independently measured functional and self-reported measures. Using the Marsbar toolbox for SPM (http://marsbar.sourceforge.net, (Brett, Anton, Valabregue, & Poline, 2002)), individual GM density values from each of the 4 DMN ROIs were extracted. These extracted GM densities were then used to test between-group differences, for correlation with the independent functional (activation-related) measures or with the independent questionnaire scores (see Figures 2 and 3).

### fMRI stimuli and experimental design

Before entering the scanner participants filled out the self-report measures and were then introduced to the MRI setup. The fMRI scans started with a resting-state of 7 minutes, in which participants were instructed to rest while keeping their eyes closed. This was followed by a visual recognition scan (1-back task in an interleaved short block design (9 s blocks with 6 s interleaved blank fixations)). Visual stimuli included stationary line drawings of faces, buildings, common man-made objects, and geometric patterns (16 of each type). Nine images from the same visual category were presented in each block; each image was presented for 800 ms followed by 200 ms uniform white screen. A central red fixation point was present throughout the experiment. Each experimental condition was repeated 7 times in pseudo-random order. The participant’s task was to fixate on the central fixation dot, and report via response box whether the presented stimulus was identical to the previous stimulus or not (1 or 2 consecutive repetitions of the same image occurred in each block). Presentation® software (Neurobehavioral Systems, Inc.) was used to deliver the visual stimuli. More details about the fMRI scan that was used for the correlation analyses are elsewhere (Berkovich-Ohana, Harel, Hahamy, et al., 2016).

### fMRI data analyses

The fMRI data analyses are detailed elsewhere (Berkovich-Ohana, Harel, Hahamy-Dubossarsky, et al., 2016), and we provide here a general description.

#### Preprocessing

fMRI data were analyzed using “BrainVoyager” software package (Brain Innovation, Maastricht, Netherlands) and complementary in-house software written in MATLAB (Mathworks, Natick, MA). Preprocessing of all functional scans included 3D motion correction, filtering out of low frequencies up to 2 cycles per scan (slow drift), and spatial smoothing using a Gaussian kernel with a full width at half maximum (FWHM) of 6 mm. Segments showing movement artifacts larger than 1 mm or sharp head movements were excluded from the analyses, without concatenating the remaining data. In addition to these pre-processing steps, resting-state data were temporally low pass filtered with a cut off frequency of 0.1 Hz (Cordes et al., 2000, 2001). The contribution of small head movements to the BOLD signal was identified using a scrubbing procedure (Power et al., 2011). To regress out non-neuronal contributions to the BOLD signal across the brain, the following procedures were carried out for each individual participant: Ventricle and white-matter regions of interest (ROIs) were manually defined while carefully avoiding the boundaries between tissue-types; these ROIs’ mean time courses were then extracted; the motion parameters, ventricle and white-matter time courses were removed from each voxel by linearly regressing them out. The anatomical scans were used to reconstruct the participant’s brain anatomy in Talairach coordinate system and then the preprocessed functional images were incorporated into 3D Talairach space (Talairach & Tournoux, 1988) through trilinear interpolation.

#### Fitting the visual recognition data to a GLM model

In order to define the L-Prc for each individual according to functional criteria, and for the functional regions of interest (fROI) task-related functional measures’ analysis, we fitted the visual recognition data of each individual to a GLM model and identified the deactivation peak within their L-Prc (see fROI definition below). Briefly, for the visual recognition first level (single subject) analysis, a general linear model (GLM, Friston et al., 1994) was fit to every voxel, with a regressor for each experimental condition based on box-car functions convolved with a hemodynamic response function. A hemodynamic lag of 6 s was assumed for all participants. Correction for small head motion was achieved by adding to the model six predictors of no interest corresponding to head motion in 3 translational and 3 rotational axes. Second level (multi-subject) analysis was based on a random-effects GLM (Zeger & Karim, 1991).

#### Functional regions of interest (fROI) definitions

The L-Prc fROIs were defined for each participant individually in Talairach space (Talairach & Tournoux, 1988), as all the voxels in the L-Prc that were deactivated in response to visual stimuli during the visual recognition task and that were within a spheres of 10 mm radius centered around that individual’s peak deactivated voxel. The maps were inspected individually and compared between participants to ensure that the L-Prc anatomical locations did not vary considerably across participants. We found that the inter-individual differences in the fROI locations did not exceed 15 functional voxels (mean of 6 voxels – Berkovich-Ohana et al., 2016).

#### Functional measures used for correlation analyses with anatomy

We used the L-Prc fROIs (see previous section) for the correlation analyses with the L-Prc ROIs (used for GM density analysis, defined in section 3.4). The ROI and fROI were not identical as each was based on different normalization procedures. The anatomical VBM normalization relied on warping the spatially smoothed GM segmented brain onto a spatially smoothed average template following DARTEL normalization procedure (Ashburner, 2007) in MNI space, while the functional normalization relied on non-linear Talairach normalization (Talairach & Tournoux, 1988) of the non-smoothed whole brain (GM, WM and other tissues). Since these methods are very different and lead to different warping of space, we opted to base our analysis on substantiated anatomical and functional regions of interest that are highly proximal within the L-Prc. For the L-Prc fROI, we used two measures, the first was the peak-to-peak activations during the visual recognition 1-back task, and the second was the BOLD variance during resting-state. To compute the task peak-to-peak activations measure for each participant and a given fROI, we extracted the fROI’s mean time course (across all its voxels) and calculated the average event-related average time course (across all visual experimental blocks). Baseline was determined as the averaged signal amplitude of all the experimental time points that preceded stimulus (block) onset by one or two TRs. The peak-to-peak differences between the average event-related average time course and baseline (fixation intervals) were calculated for each fROI and participant separately as the difference between the maximum and minimum activation values (in percent signal change (PSC) units) within the 0-9 s time window after stimulus onset. The resting state BOLD variance measure reflects the average fluctuations in the signal amplitude during spontaneous resting-state activity. For each participant and each fROI the average resting state fROI time-course was extracted (x(t)=x_1_,x_2_…x_180_) and then converted to PSC (relative to the time course mean 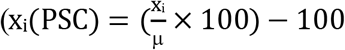, where μ is the time-course mean). The variance of that entire time-course was calculated for each participant as 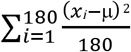. Data of one MM participant were excluded from the analysis due to values exceeding ±2 standard deviations from the group average.

### Factor analysis

The factor analysis incorporated the following steps: An unrestricted principal components (PC) solution, using a loading of 0.55 as criterion for marking those items loading on each factor, from which the number of factors could be determined by scree test, followed by a restricted factor analysis; factor solutions were then rotated orthogonally using Varimax (all analyses being conducted on SPSS). With present sample sizes (*n* = 33) the ratio of *n* to the number of items entering into the analysis is greater than 5 : 1 which is satisfactory.

## Results

Following inconsistent earlier findings about meditation experience and the DMN neuroanatomy (reviewed by Fox et al., 2014), our first goal was to examine whether meditation experience would be correlated with the gray matter (GM) structure of the DMN, and whether any group differences between long-term mindfulness meditation (MM) practitioners and controls over DMN regions could be found.

Voxel-based morphometry analysis in each of the 4 DMN ROIs, revealed a significant MM vs. control group difference only in the L-Prc (the most significant peak at MNI coordinates: −21, −64, 16 (BA 7), *p* = .013, family wise error (FWE) corrected) (Figure 1A). In an additional, more refined, VBM analysis, this region was also found to show significant correlation between GM density and meditation experience (by years) (peak at MNI coordinates: −21, −64, 16, p < .005, FWE corrected). We extracted GM peak values from this focus to verify that the correlation is not driven by outliers, and to examine whether the correlation would stay significant when only taking into account the MM group. As can be seen in Figure 1B, the overall correlation across both groups (in gray, [*r* = −.703, *n* = 36, *t*(34) = −5.771, *p* < 0.0001]) is not driven by outliers. This remained statistically significant following Holm-Bonferroni corrections for multiple comparisons. The correlation remains significant when based only on the MM group (Figure 1B, depicted in red, [*r* = −0.526, *n* = 18, *t*(16) = −2.478, *p* = 0.0247]). However, after Holm-Bonferroni corrections for multiple comparisons, this correlation for the MM group is no longer significant. Further whole-brain analysis examining whether these structural meditation-related effects may extend beyond the DMN did not reveal any other significant clusters.

These results indicate that the neuroanatomy of the L-Prc but not of other ROIs reflects long-term MM practice experience. We thus went on to investigate structure – function and structure – self-report relationships, focusing on this region, and for that purpose we extracted GM density from the L-Prc ROI (see Methods).

We first hypothesized a general structure-function correlation, and tested it over the two groups together, for two functional measures, but this yielded no significant correlations (Table 1). However, when testing the same correlation separately for each group, we found only for the MM practitioners that GM density was positively and significantly correlated with the functional measures (visual task peak-to-peak measure [r = .605, n = 15, t (13) = 2.74, p = .017], and resting-state functional measure [r = .552, n = 15, t (13) = 2.39, p = .033]) (see Figure 2 and Table 1). These structure-function correlations within the MM group remained statistically significant following Holm-Bonferroni corrections for multiple comparisons. Assessing whether the correlations of the MM and the control groups were significantly different, we found the inter-group correlation coefficient difference to be marginally significant [two-tailed *z* = 1.88, *p* = .060, same statistical results for both the resting state variance and task peak-to-peak activity analyses]. We then analyzed both groups together, but this analysis did not yield any significant structure-function results.

Finally, we examined the relationship between the L-Prc’s neuroanatomical structure and individual self-reported measures by correlating GM density values with PANAS scores or with TAS scores, first over both groups together, and then separately for each group, as before (Figure 3 and Table 1, for group differences see Table 2). Comparing the positive affect scores with GM density across all the participants, we found a significant negative correlation [*r* = −.522, *n* = 33, *t* (31) = −3.41, *p* = .002], which survived the Holm-Bonferroni correction. No significant correlations were found with either negative affect or TAS scores. When examining these correlations in the two groups separately, we found that only the MM group correlation was significant (Figure 3), indicating that the all-participant correlation was driven by the MM group. We computed correlation matrices to investigate how all the measures inter-relate, both across the whole group and within each group (Table 3), as well as a factor analysis. The initial PC analysis revealed 2 factors with eigenvalues greater than unity, which accounted for 60.34% of the variance, and were employed in the next phase of the analysis. An orthogonal Varimax rotation revealed simple structure as follows: the first factor (38% of the variance) was identified as with positive and negative affect (PA, NA), with L-Prc GM density and fMRI peak-to-peak items loading at 0.55 or better. The second factor (22.34% of the variance) was identified as absorption (TAS) with fMRI variance loading at 0.55 or better. Thus, when one incorporates all relevant factors in the analysis, as seen from the factor analysis, it would seem that structural and functional factors are partly distinct: while emotional components lean on L-Prc structure and fMRI peak-to-peak task-related measure, the absorption is related to L-Prc fMRI resting state variance.

**Table 2.** Questionnaires scores (mean ± SD) and statistical comparisons between groups (1-way ANOVA). MM – Mindfulness meditators (*n* = 16); Controls (*n* = 17); PA – positive affect; NA – negative affect; TAS – Tellegen’s absorption scale. * *p* < .05.

| Questionnaire | MM | Control | *F-contrast* |
| --- | --- | --- | --- |
| PA | 39.1 ± 5.0 | 36.2 ± 4.0 | 3.25 |
| NA | 19.1 ± 5.8 | 20.0 ± 5.3 | .21 |
| TAS | 22.0 ± 7.6 | 16.1 ± 6.6 | 5.60* |

**Table 3.** Pearson correlations matrix between all measures: L-Prc ROI neuroanatomical gray matter (GM) density (structure) and fMRI BOLD functional measures (Z-transformed variance during resting-state (L-Prc var), and peak-to-peak activation values during a visual task (L-Prc p2p) or self-reported scores on questionnaires (PANAS and TAS – see Methods). MM – Mindfulness meditators (n = 16); Controls (n = 17); * p < .05; ** p < .005.

|  |  | **L-Prc**<br>**GM** | **PA** | **NA** | **TAS** | **L-Prc**<br>**p2p** |
| --- | --- | --- | --- | --- | --- | --- |
| **All (*n* = 33)** | **PA** | -.522** |  |  |  |  |
|  | **NA** | .158 | -.037 |  |  |  |
|  | **TAS** | -.305 | .379* | .118 |  |  |
|  | **L-Prc p2p** | .294 | -.433* | .160 | .034 |  |
|  | **L-Prc var** | .270 | -.048 | -.015 | -.440* | .208 |
| **MM (*n* = 15)** | **PA** | -.555* |  |  |  |  |
|  | **NA** | .028 | .246 |  |  |  |
|  | **TAS** | -.426 | .497 | -.395 |  |  |
|  | **L-Prc p2p** | .605* | -.247 | .053 | -.015 |  |
|  | **L-Prc var** | .552* | .040 | .078 | -.394 | .203 |
| **C (*n* = 17)** | **PA** | -.371 |  |  |  |  |
|  | **NA** | .425 | -.340 |  |  |  |
|  | **TAS** | .169 | .007 | .706** |  |  |
|  | **L-Prc p2p** | -.025 | -.507 | .159 | .217 |  |
|  | **L-Prc var** | -.102 | .082 | -.209 | -.336 | .041 |

## Discussion

Here we investigated whether during adulthood brain structure is modulated by cognitive mental training in the form of mindfulness meditation (MM) training, and how that relates to brain function and self-reports. We found that duration of MM practice was predictive of GM density in the left precuneus (L-Prc), part of the DMN - a network associated with inner thought related processes such as episodic memory and mind wandering (e.g. Addis, Wong, & Schacter, 2007; Bar, 2007; Mason et al., 2007), and part of the self-reference network (Kim, 2012; Northoff et al., 2006; Northoff, 2016). We further found that GM density in the L-Prc was positively correlated to activation during task or variance during rest.

Variance measures were previously used as a metric to estimate the amplitude of spontaneous (resting state) activity fluctuations - i.e. they provide an indirect measure of how high are the slow transient activations that appear during resting state (Bianciardi et al., 2009; Davis, Jovicich, Iacovella, & Hasson, 2013; McAvoy et al., 2008). As was demonstrated in numerous previous studies, these activity fluctuations do not emerge randomly in isolated voxels, but show coherency across cortical networks. This coherency in the fluctuations has been termed “functional connectivity”. However, it is important to draw a distinction between the amplitude of the local fluctuations (measured through the variance) and the level of correlation between these local fluctuations (measured through functional connectivity). According to the “spontaneous trait reactivation” (STR) hypothesis (Harmelech & Malach, 2013), the connectivity patterns of the spontaneous fluctuations recapitulate training-induced individual differences, which could be observed during task performance, i.e. habitual patterns of brain activations should be correlated to unique changes in spontaneous patterns of activations that are generated during rest. Importantly, our previous functional analyses in the same cohort of participants reported here supported the STR hypothesis, revealing function – variance correlation (see the “relative activation index” analyses, Berkovich-Ohana, Harel, Hahamy, Arieli, & Malach, 2016). At the same time, variance was not significantly correlated to functional connectivity measures (which rely on measuring network correlations) for this cohort, supporting the proposition that these measures differ in relation to the brain activity they convey. The lower variance in the meditators compared to the controls, as well as their correlation with task activity in the same regions, were previously conjectured as reflecting a long-term reduction of DMN activity in MM practitioners, i.e. reflecting a long-term reduction of internal, self-related processing. This interpretation is further strengthened by the significant structure – function – expertise correlation reported here. However, as our results build on no randomization and only a single time point of measurement, it only provides suggestive correlations, and we cannot rule out, at this point, the alternative interpretation - that both anatomical changes as well as the functional measures reflect a-priory personality trait that leads specific individuals to engage in meditation to a specific extent. Resolving this question will necessitate a long-term meditation intervention randomized controlled experiment that is beyond the scope of our current study.

Brain plasticity during adulthood and its causes and influences is a field of immense interest (e.g. Kehayas & Holtmaat, 2017; McEwen, 2016). The reduced GM density we found in the L-Prc associated with prolonged MM practice is in line with Kang and colleagues (2013) showing reduced GM thickness in left posterior cingulate (BA 31), left precuneus (BA 7) and cuneus (BA 17) in long-term meditators (3.4 ± 2.3 yrs) compared to controls. Yet, other studies show increased PCC GM thickness following a short 8-week mindfulness intervention (Hölzel, Carmody, et al., 2011; Yang et al., 2019; and see review by Fox et al., 2014). One possibility to settle this discrepancy is by suggesting that GM density in the precuneus is modulated by MM practice according to an inverted U-shape, so that there is an initial rise in GM thickness following the initial phase of MM practice (several weeks, as in MBSR) and then GM starts to reduce with continued MM practice (as in our data, where practitioners accumulated years of practice). In line with this suggestion, theoretical models and animal work predict that regional volumes may rapidly expand during learning and then partially renormalize, despite continued training (Lövdén et al., 2013), and in humans GM volumes follow inverted U-shaped developmental curves during childhood (Lenroot & Giedd, 2006).

Here we report that structure of the L-Prc, an amodal area, is associated with prolonged cognitive mental practice within the MM practitioners (see Results and Figure 2). The GM reduction we found in L-Prc in the MM compared to the control group was not only anatomically close to our fMRI previously reported results (Berkovich-Ohana, Harel, Hahamy, et al., 2016), but also proximal to foci reported in other meditation studies, including earlier neurofeedback studies by Brewer and colleagues (Brewer et al., 2011; Garrison et al., 2013) (cf. L-precuneus MNI coordinates in our study (−8, −59, 27) and the Brewer study (−6, −60, 18)), as well as Pagnoni’s (2012) paper on the ventral posteromedial cortex (vPMC) skeweness (MNI coordinates: −6, −56, 22). These earlier studies suggest that activity in this region is supporting “getting caught up in the experience”, while reduction in its activity supports “undistracted” and “effortless awareness”, as well as “contentment” (Garrison et al., 2013). Hence, the L-Prc region is suggested to support the experiences of identifying with or being attached to attributes of ourselves (Brewer et al., 2013). Thus, our results support the possibility that brain structure and function outside modality-related cortex could be co-modulated during adulthood in association with mental training.

The finding of a significant association between structure and function only in the MM group and not in controls is not straightforward. One possible and rather intuitive explanation is that there are inter-individual differences in the processes associated with the precuneus, e.g. self-referential attribution/attachment to internal experiences (Garrison et al., 2013; Northoff, 2016), and once people start to meditate both structure and function become more tuned to these processes (Brewer & Garrison, 2014; Brewer et al., 2013). With meditation, over years of practice, mental control improves, and structure and function become tuned to these mental functions and their relation is amplified as the years go by. Such an explanation is supported by previous reports of enhanced stability in brain activity in meditators compared to controls (Lutz et al., 2009). However, there might be another less straightforward explanation for the lack of correlation in controls. Hypothetically, there is a possibility that in the typical (meditation naive) brain, there would be certain above-baseline average activity and a certain average structure (GM density), with inter-individual variability in these two measures around these average values. In such a case, these two values would not be correlated in non-meditating individuals. However, if meditation experience acts as a reducer of function and a thinner of structure, then it could be that meditation not only reduces the inter-individual variability in both of these measures, but also at the same time reduces the average values of each of these measures. In such a scenario, correlation between these two measures could be found as a consequence of these co-occurring synchronized reductions, and not necessarily because they are directly related. In our case of MM practitioners, the meditation experience spanned a wide range of durations (8 - 35 years), which according to such a hypothetical scenario may give rise to such a correlation.

The relationship between brain structure,function and behavior is not always clear from cross-sectional studies, and may be brain region dependent. This is especially the case for the aging brain: a growing literature on healthy aging shows a positive structure-function relation in task-related PFC areas, with positive correlation to task performance (reviewed by Maillet & Rajah, 2013), in support of the dedifferentiation model of age-related changes in brain function (postulating age related reductions in the signal-to-noise ratio and regional specialization of function (Li, Lindenberger, & Sikström, 2001)). In contrast, in pathological aging and dementia a positive function-structure relationship was observed in PFC task related areas (reviewed by Maillet & Rajah, 2013), in support of the compensatory neural plasticity hypothesis (postulating over-activations in task-related regions, to compensate for declining neural efficiency (Cabeza, Anderson, Locantore, & McIntosh, 2002)). Thus, different and opposing structure-function mechanisms may be at play in the aging brain. Conflicting evidence is also true for the healthy adult brain: on the one hand, multiple studies suggest that more neural machinery dedicated to specific perceptual/cognitive functions could provide more precise coding for that behavior, thus a positive structure-performance relationship (Gilaie-Dotan et al., 2014; Grubb, Tymula, Gilaie-Dotan, Glimcher, & Levy, 2016; Schwarzkopf et al., 2011). For example, a recent study (Schwarzkopf et al., 2011) suggests that V1 surface area 3-fold variations in neurotypical adults (Adams, Sincich, & Horton, 2007; Andrews, Halpern, & Purves, 1997) may be related to differences in subjective size perception. On the other hand, Maguire and colleagues (2000), report that within the same individuals two anatomically adjacent locations in the hippocampus are modulated in opposite directions by navigation experience, where years of experience as a taxi driver positively correlate with anatomy of the right posterior hippocampus, but negatively with that of the anterior hippocampus. It is also an open question whether the same mechanisms underlie the adult and aging brain (Lövdén, Wenger, Mårtensson, Lindenberger, & Bäckman, 2013). At the same time, cross-sectional studies showed that people with high levels of expertise, generally show greater GM volume compared to non-experts in parts of the brain crucial for that relevant expertise. For instance, professional musicians, showed greater GM volume compared to non-musicians in brain regions assumed crucial for fine motor control and auditory processing (Gaser & Schlaug, 2003; Schneider et al., 2002). While such cross sectional studies suggested that training had induced the GM inter-individual differences, i.e. more neural machinery provides more precise coding for a specific skill, a plausible alternative explanation is that differential GM volume would be the cause, not the consequence, of long-term practice. In order to establish whether training itself had been the cause of GM volume differences, GM volume was to be investigated before and after training in a longitudinal fashion. This limitation was addressed by a few longitudinal training studies, where accumulated evidence supports the notion that a positive function-structure-performance correlation was causally driven by training effect. This includes a wide variety of experimental paradigms, such as juggling (Draganski et al., 2004) (Driemeyer, Boyke, Gaser, Büchel, & May, 2008), golfing (Bezzola et al., 2011; Boyke et al., 2008), and musical training (Hyde et al., 2009). Based on the above, our cross-sectional results of positive structure-function correlation cannot suggest causality, and the question of whether it stems from MM training remains to be explored in a longitudinal training study.

In our study we found a negative correlation between both L-Prc structure and function and self-reported positive affect scores supporting the previously suggested link between enhanced happiness and reduced self-referential processing (Dambrun & Ricard, 2011) or mind-wandering (Fell, 2012; Killingsworth & Gilbert, 2010), both experiences associated with DMN activity or its sub-systems (Andrews-Hanna, Reidler, Sepulcre, Poulin, & Buckner, 2010; Northoff et al., 2006; Schacter et al., 2007). Interestingly, other studies link a reduction in precuneus GM density to reduced pain sensitivity (Emerson et al., 2014), and enhanced present-moment awareness and trait-mindfulness (Lu et al., 2014). Another study reports that the spontaneous tendency to recall memories from a first-person perspective is positively correlated with the precuneus GM volume among two independent datasets (Freton et al., 2014). This is aligned with accumulating evidence showing that self-relevant versus neutral stimuli activates the precuneus (Herwig et al., 2010; Lutz et al., 2016), while mindful self-awareness decreases it (Lutz et al., 2016). Moreover, mental process of directing attention and awareness to emotions and bodily feelings, notably without the conscious intention to regulate emotions, as done in mindfulness practice – has the ability to attenuate emotional arousal related brain activation (Herwig et al., 2010). Indeed, the precuneus in long-term meditators compared to novices shows decreased functional connectivity to dorso-medial prefrontal cortex (the frontal node of the DMN) when facing both self-criticism and self-praise, illustrating the mechanisms of mindful emotion regulation as differences in self-related emotional processes (Lutz et al., 2016). Additionally, the precuneus exhibits enhanced functional connectivity patterns with subcortical affective networks following meditation training, and the connectivity between the precuneus and pons predicted changes in affective processing after meditation training, suggesting that meditation promotes self-referential affective regulation based on increased regulatory influence of the pons on PCC/precuneus (Shao et al., 2016). The above studies provide evidence that mindfulness practice fosters an even-minded mental state of ‘equanimity’, ultimately developed towards all experiences, regardless of their affective valence (Desbordes et al., 2015). Accordingly, mindfulness meditation leads to less differentiated affective processing of emotionally valenced stimuli, as well as reduced general affective reactivity and arousal (Farb et al.,2010; Goldin and Gross, 2010).

In line with the above logic suggesting that more neural machinery provides more precise coding for a specific skill, if MM practice reduces self-related focus and processing, and the precuneus supports self-referential processing, then the more MM is practiced, the less self-referential machinery and processing should take place. And indeed, here we found that not only the L-Prc’s GM was reduced in MM practitioners as a function of practice duration, but that also brain activity in these practitioners was proportionally reduced. While this logic might not provide an across the board explanation to differences or changes in positive affect, it can suggest that structure and function both have significant roles, and might both influence or be associated with positive affect.

It should be noted that our study has several limitations. First, the sample size in this study was relatively small, which renders the results as preliminary and exploratory, requiring replication in a larger sample. Second, the study we conducted was not longitudinal, but cross-sectional and self-selective, and thus limits the causal inference that could be drawn from it. Hence, the results did not show that meditation practice caused reduction in GM of the L-Prc, but rather suggests a correlation between GM density and accumulated meditation practice, which is evident at a single time-point with a self-selecting cross-sectional sample of meditators. While our approach enables studying expert MM practitioners, with accumulated thousands of hours of practice, longitudinal studies with random assignment to groups may address this shortcoming and provide more direct insights to the question of causality.

Nevertheless, our results shed light as to how structure and function are both tightly linked and associated with mental practice and subjective affective reports. The finding that both structure and function of the L-Prc are correlated, as well as associated with a specific mental practice can shed light on the mechanisms and processes that this area is involved in and tuned to, specifically emphasizing the role of this area as a target of meditation training, as previously suggested (Brewer & Garrison, 2014).

## Acknowledgements

The study was funded by the Helen and Kimmel Award for innovative Research, the EU (FP7 VERE), the EU - Human Brain Project and the ISF-ICORE grants to R. Malach, the Teva Pharmaceutical Industries LTD fellowship to A. Berkovich-Ohana, and the ISF Individual Research Grants to S. Gilaie-Dotan (no. 1485/18). Dr. E. Furman-Haran holds the Calin and Elaine Rovinescu Research Fellow Chair for Brain Research.

